# Tuberculosis causes highly conserved metabolic changes in human patients, mycobacteria-infected mice and zebrafish larvae

**DOI:** 10.1101/2020.05.25.114470

**Authors:** Yi Ding, Robert Jan Raterink, Rubén Marín-Juez, Wouter J. Veneman, Koen Egbers, Susan van den Eeden, Mariëlle C. Haks, Simone A. Joosten, Tom H.M. Ottenhoff, Amy C. Harms, A. Alia, Thomas Hankemeier, Herman P. Spaink

## Abstract

Tuberculosis is a highly infectious and potentially fatal disease accompanied by wasting symptoms, which cause severe metabolic changes in infected people. In this study we have compared the effect of mycobacteria infection on the level of metabolites in blood of humans and mice and whole zebrafish larvae using one highly standardized mass spectrometry pipeline, ensuring technical comparability of the results. Quantification of a range of circulating small amines showed that the levels of the majority of these compounds were significantly decreased in all three groups of infected organisms. Ten of these metabolites were common between the three different organisms comprising: methionine, asparagine, cysteine, threonine, serine, tryptophan, leucine, citrulline, ethanolamine and phenylalanine. The metabolomic changes of zebrafish larvae after infection were confirmed by nuclear magnetic resonance spectroscopy. Our study identified common biomarkers for tuberculosis disease in humans, mice and zebrafish, showing across species conservation of metabolic reprogramming processes as a result of disease. Apparently, the mechanisms underlying these processes are independent of environmental, developmental and vertebrate evolutionary factors. The zebrafish larval model is highly suited to further investigate the mechanism of metabolic reprogramming and the connection with wasting syndrome due to infection by mycobacteria.

## Introduction

Infection with *Mycobacterium tuberculosis (Mtb)* can result in tuberculosis (TB) that currently is newly affecting over 10 million people per year and leading to 1.6 million deaths annually^1^. However, the majority of people that have been infected with *Mtb*, estimated to be in the order of 2 billion, do not experience any symptoms in the presence of positive immunological (blood or skin) test results. In patients diagnosed with TB, the disease is uniformly characterized by a metabolic wasting syndrome that leads to heavy weight loss that concurs with a strong reduction of muscle mass. TB has therefore historically earned the name of “consumption”.

Novel tools for control of TB should include not only better treatments, including attention to the deleterious effects of wasting syndrome, but also needs to include cost-effective assays for robust and sensitive diagnosis of TB^2^. Recent metabolomics studies have identified valuable diagnostic metabolic blood markers that can discriminate TB patients from healthy individuals^3–13^. This appeared to be applicable to very different population groups, ranging from trans-Africa, China, Georgia, and Indonesia, and therefore highlighting that blood metabolites are robust and validating biomarkers for TB progression. These studies used mostly mass spectrometry (MS) and in some cases nuclear magnetic resonance (NMR) spectrometry^4^. Compared to MS, NMR offers the advantages of high reproducibility, minimal sample preparation, fast measurement, non-invasive, non-selectivity, non-destructive such that samples also could be stored and used for other detections. The limitation of NMR is its relatively low sensitivity, which however can be improved by using higher magnetic fields (frequencies commonly used range between 300 and 700 MHz), and large numbers of samples^14,15^. MS studies of metabolites from blood samples showed that many amino acids, such as histidine, cysteine, glutamine, tryptophan, citrulline, and creatine were decreased in TB patients from South Africa^3^. A few metabolites were increased in abundance including kynurenine, phenylalanine and pyroglutamine in the active TB group^3^. Similar patterns of metabolic changes due to TB were found in a recent MS study on blood samples from a cohort of Indonesian TB patients even if the patients suffered from a combination of TB and type II diabetes^16^. By comparing metabolic profiles of TB patients from patients with other respiratory diseases, Weiner et al. identified metabolic profiles that are specific to TB^11^. In this study it was shown that changes in various metabolites were predictive for the onset of clinical TB as early as 12 months prior to TB diagnosis in a trans-African population^11^. Using MS, Yi et al. demonstrated that histidine, arachidonic acid and biliverdin were decreased in Chinese TB patients compared to the group of healthy individuals, but could be restored to normal levels following TB cure^12^. In an MS study that compared the effects of curing patients from Gambia of an infection of *Mtb* or *M. africanum* on metabolism, it was shown that in both cases the levels of many amino acids such as histidine, ornithine, tryptophan and leucine were increasing strongly after 6 months of antibiotic treatment^7^. In a study that used NMR for analysis of blood metabolites, markers for Chinese TB patients were found, although there were major differences with the published MS studies, such as an increased plasma level of glutamine^4^.

There are only few studies investigating the metabolic profile of *Mtb* in animal models, such as cattle, guinea pigs and mice using NMR. A study with calves that were infected with three *Mtb* strains including *M.bovis, H37Rv* and *Mtb1458* through intratracheal injection and followed for 33 weeks showed that serum levels of isoleucine, leucine, tyrosine, valine were decreased in case of all three strains^17^. Using ^1^H Magic Angle Spinning NMR, Somashekar et al. revealed that alanine was significantly elevated after 15 and 60 days infection with *Mtb H37Rv* in lung tissue of challenged guinea pigs^18^. A few metabolites such as ethanolamine, glutamine and ATP were significantly decreased after 30 and 60 days of infection in serum of guinea pigs^19^. Clear separation between control and *Mtb*-infected groups was also observed in organs (lung, liver, spleen) and serum samples of mice^20^. A majority of these metabolites including tyrosine, glutamine, leucine, isoleucine were increased in lung, liver, spleen and serum^20^. Although these NMR studies identified metabolic markers for TB, there was little correlation with the markers identified in human studies using MS.

Zebrafish has been shown to be an excellent model for the study of TB^21–25^. Because zebrafish have large amounts of offspring at one time and the availability of high throughput automated infection methods^26^, many zebrafish larvae can be tested for TB in a short time. The test system is independent of food intake and adaptive immune responses that only start later during development. In order to validate our platform for identification of TB biomarkers, we also performed metabolomic studies in zebrafish larvae. It is an important model for TB research based on the infection of zebrafish larvae with *Mycobacterium marinum (M.marinum)*. It has been shown that this infection leads already after several days to many inflammatory responses and state that are highly comparable to progression of TB in humans^27,28^. The sensitivity of our mass spectrometry platform has been shown to be sufficient for the analysis of single zebrafish larva for the detection of large set of small metabolites^29^.

Here, we have compared the effect of TB infection on the level of metabolites in blood of humans and mice and whole zebrafish larvae using one highly standardized mass spectrometry pipeline, ensuring technical comparability of the results. Subsequently we used the zebrafish model to further investigate the metabolic changes after infection using NMR analyses based on a large number of specimens. Our results show a common conservation of metabolic reprogramming processes during TB infection and disease in humans, mice and zebrafish. The NMR analysis of zebrafish larvae confirmed the MS data and also identified new markers for infection including identifying general effects on metabolism such as a change of glucose levels.

## Results

### Metabolite profiles of TB patients measured by mass spectrometry

To measure metabolite profiles in TB patients compared to healthy people, 20 blood samples from each group were analysed by MS. Using a highly standardized platform we could measure 59 small amine-containing compounds. The normalized data sets showed a significant separation of the metabolite profiles of the patient and control group using Partial Least Squares Discriminant Analysis (PLSDA) (Fig.1A). Using a false discovery rate cut-off value of 0.05 we could classify 31 of the 59 identified small amine-containing compounds as associated with the TB disease state (Fig.1B, Table S2). There were 26 metabolites showing significantly lower abundance in patients compared to healthy individuals (Fig.1B). Only a few metabolites, namely putrescine, methionine sulfoxide, aspartic acid and glutamic acid, were increased in patients compared to controls (Fig.1B). Considering the relatively large number of metabolites of which the levels were significantly affected, the data suggest a drastic metabolic reprogramming in TB patients. For 8 of these metabolites it was previously shown that they were biomarkers for TB, namely methionine, glutamine, threonine, tryptophan, histidine, citrulline, cysteine and homoserine^3,11,16^. Antibiotic treatment for 6 weeks did not have a major effect on separation of the metabolic profiles of TB patients from the control group although minor effects were observed (Fig.S1). Some markers appeared to be responding in an anti-correlated fashion compared to published TB cohorts studies, such as lowered level of phenylalanine (Fig. 1B) that was shown to be increased in patients in the study of Weiner et al, 2012^3^ and Vrieling et al, 2019^13^. However, in a study of a TB patient cohort from South Africa, also a decreased level of phenylalanine was reported compared to the control group^30^.

**Fig.1.**
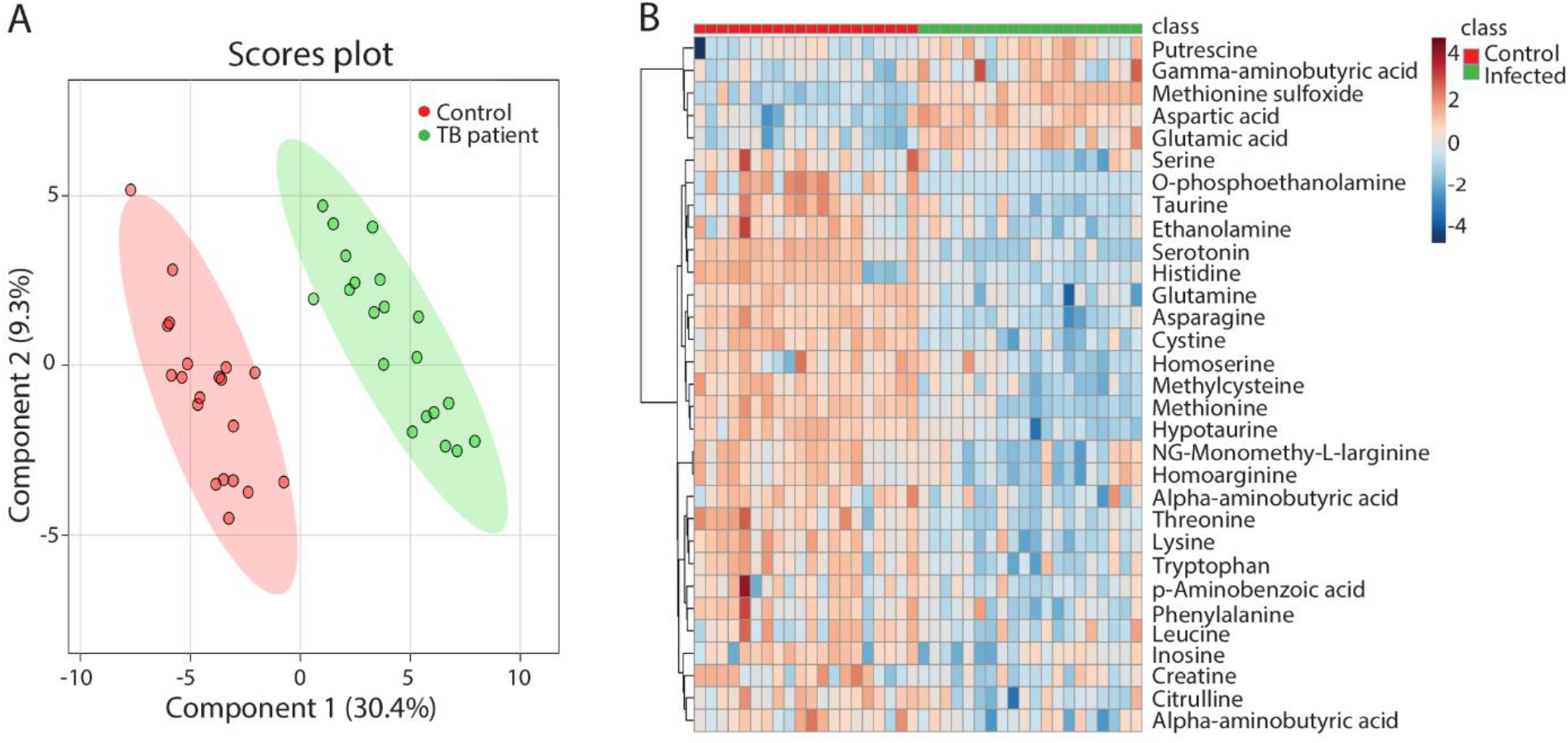
Partial least squares discriminant analysis and heat map of metabolomic profiles from patients with active TB and healthy controls. **A**. Analysis of blood of the healthy control group (Control), patients with active TB disease (TB patients), n=20. **B**. Heat map of the 31 metabolites which have a significantly different concentration in TB infected patients and the healthy control group in a T-test with *p*<0.05.

### Metabolite profiles of *Mtb* infected mice measured by mass spectrometry

Metabolic profiles of the blood of *Mtb*-infected C57BL/6 mice and an uninfected control group were measured by MS to identify and compare profiles in a rodent model of TB. PLSDA score plots of the 40 identified metabolites showed clear differences between the control group and the *Mtb*-infected mice, indicating there were significant alterations in the metabolism after infection (Fig.2A). The data showed significant differences and ratios of metabolite quantities for 31 of the 40 identified molecules between infected mice and controls by applying a t-test at a significant threshold of 0.05 (Fig.2B, Table S3). The relative abundances of all the significantly identified metabolites were reduced after infection (Fig.2B). Of these signature metabolites 14 were also lower in human TB patients compared to controls (Fig.3A). In contrast to the human data set, there was no increase of any metabolite after infection.

**Fig.2.**
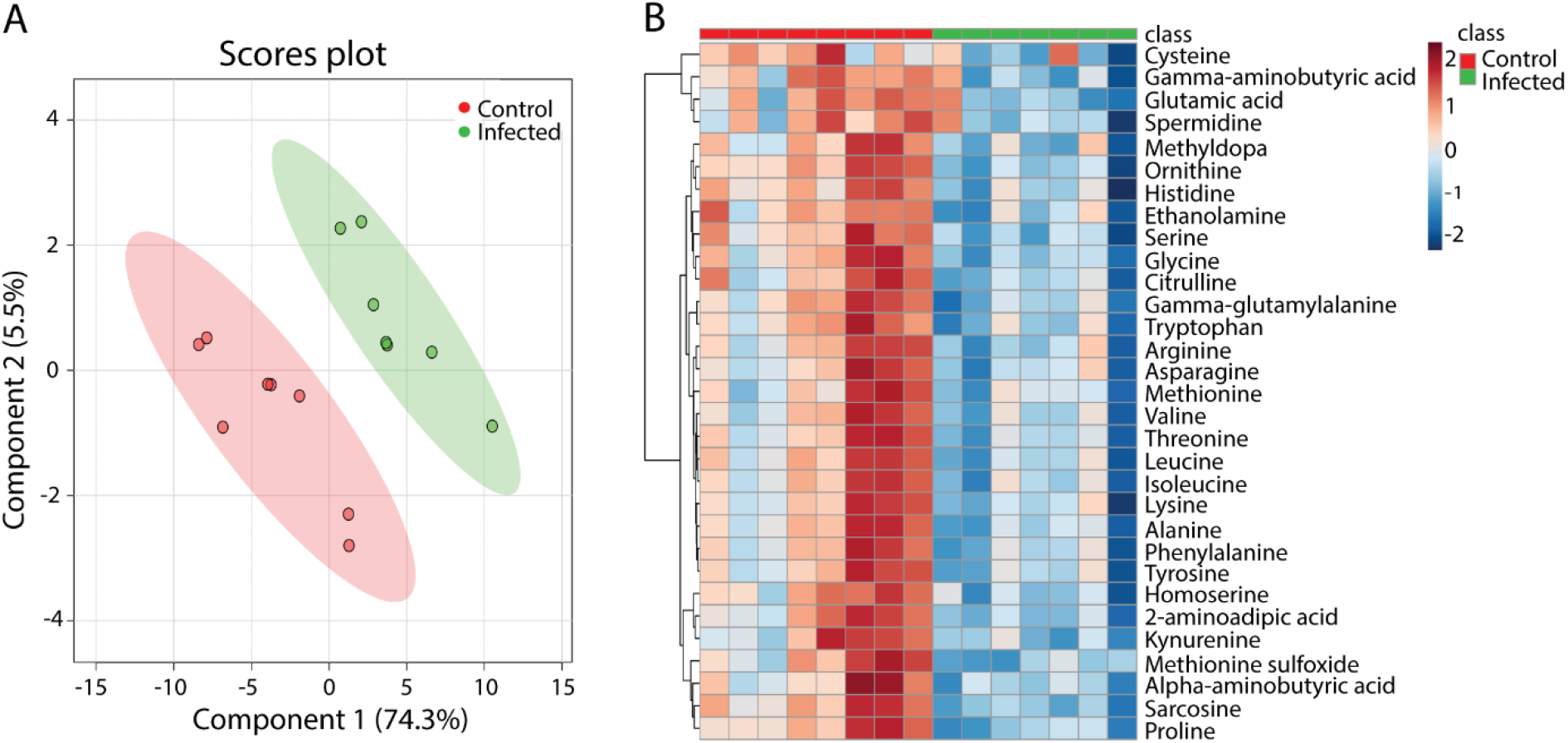
Partial least squares discriminant analysis and heat map of metabolomic profiles from mice with or without *Mtb* infection. **A**. PLSDA analysis of blood of the wild type mouse strain C57BL/6 after 8 weeks of nasal infection with *M.tuberculosis* H37Rv, n=8. **B**. Heat map of the 31 metabolites which have a significantly different concentration in *Mtb*-infected and uninfected mice in a T-test with *p*<0.05.

**Fig.3.**
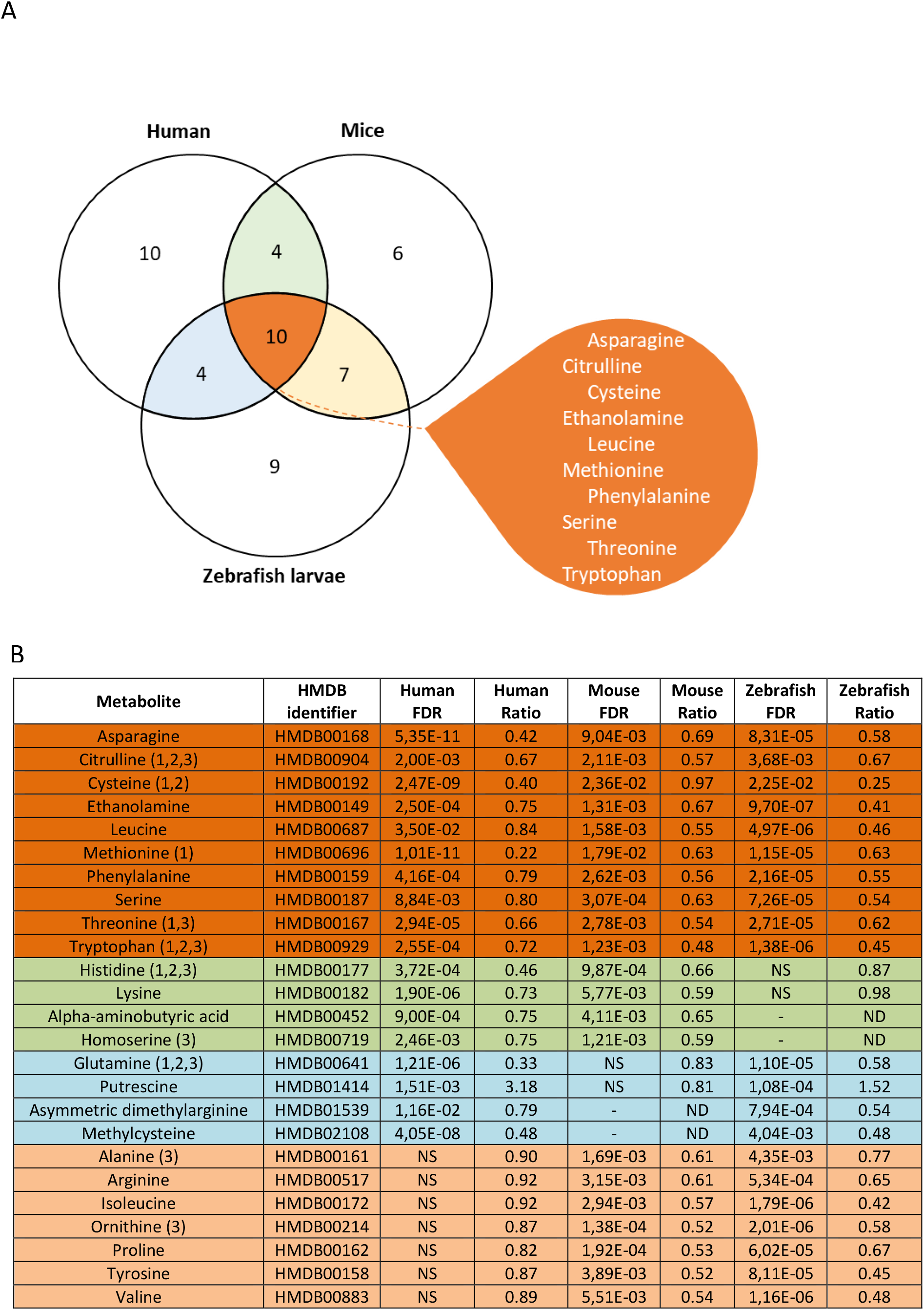
Common metabolic responses in TB patients, infected mice and zebrafish. **A.** A Venn diagram is shown of the common metabolites that respond significantly towards infection with mycobacteria in patients with TB disease, infected mice and zebrafish larvae. Anti-correlated values are not included in the overlap of the Venn diagram. **B**. A table shows the ratio of common metabolites in humans, mice and zebrafish larvae. Abbreviations: ND, non-detectable; NS, not significant. (1): Metabolites that have been reported to have significantly different levels between patients with TB as compared to healthy controls (Weiner et al, 2012). (2): Metabolites of which the level have been reported to be predictive for the development of TB in African patients (Weiner et al, 2018). (3) Metabolites that are significantly decreased in TB patients (Vrieling et al, 2019)

### Metabolic profiles of *M.marinum* infected zebrafish larvae measured by mass spectrometry

In order to test whether the metabolic responses observed in humans and mice could be translated to results in zebrafish, larvae infected for 5 days with *M.marinum* strain E11 were studied by MS. Metabolomic profiles of the 44 identified metabolites of complete infected and uninfected larvae could be clearly discriminated using PLSDA (Fig.4A). A set of 35 out of 44 identified small amine-containing metabolites was significantly different with an FDR cut-off value of 0.05 (Fig.3B, Table S4). The majority of the 30 metabolites was significantly decreased in the infected zebrafish larvae (Fig. 3B, Table S4). Five metabolites, namely glutathione, putrescine, beta-alanine, 5-hydroxylysine and O-phospoethanolamine, were increased in the infected group compared to the control group (Fig.4B). There was an overlap of 17 metabolites with the TB signature metabolites observed in mice and of 14 metabolites with the TB signature metabolites observed in humans (Fig.3A).

**Fig.4.**
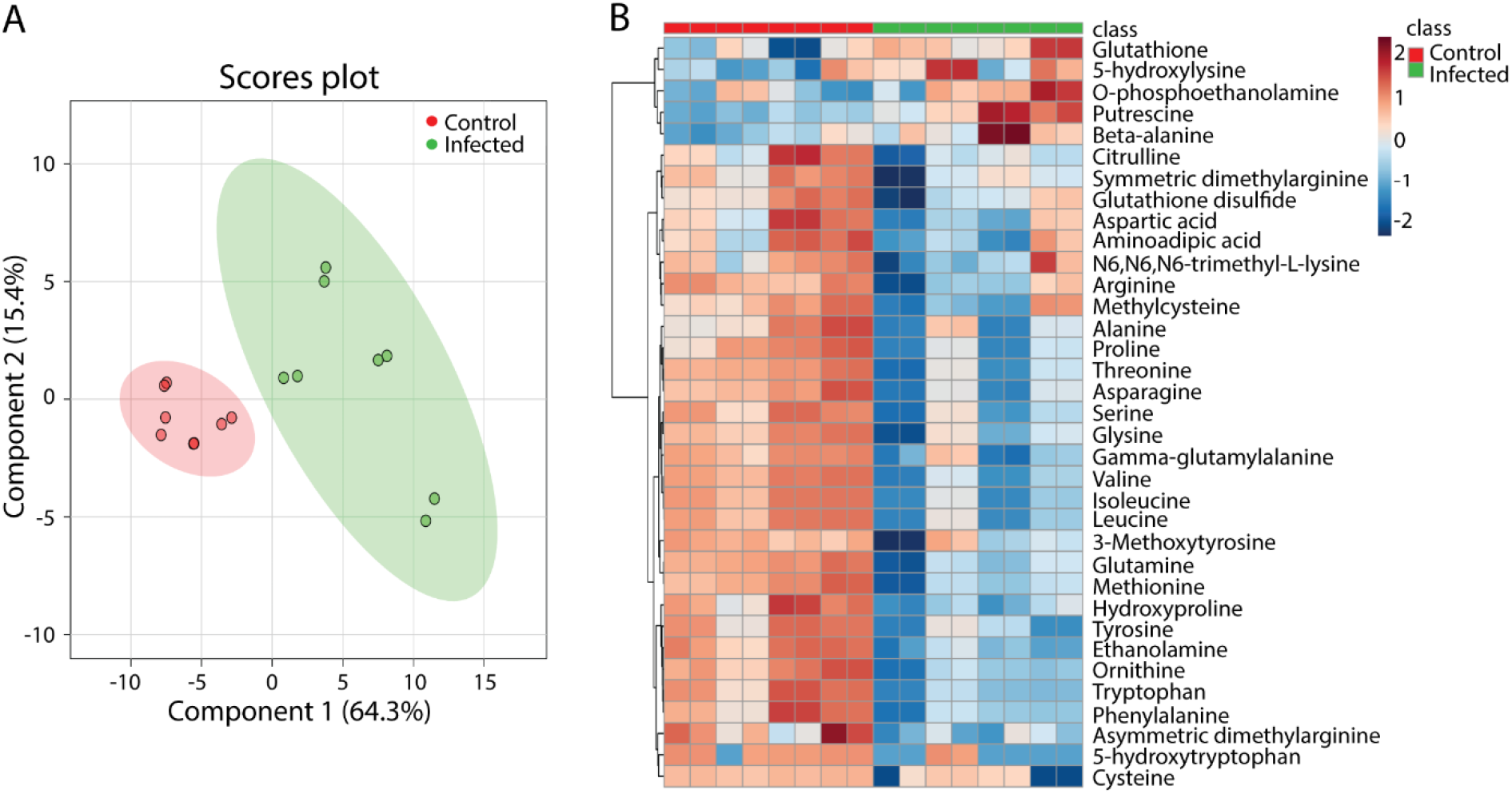
Partial least squares discriminant analysis and heat map of metabolomic profiles from *M.marinum-infected* and uninfected zebrafish larvae. **A**. Analysis of entire zebrafish larvae uninfected (Control) or after infection with *M.marinum* strain E11 in the yolk at 2 hours post fertilization (Infected). All the larvae were collected at 5 days post fertilization and then measured with mass spectrometry. Each dot represents one larva, n=8. **B**. Heat map of the 35 metabolites which have a significantly different concentration in zebrafish larvae in the infected and the control group.

### A core set of metabolites are markers for tuberculosis in humans, mice and zebrafish

Interestingly and consistently, the three metabolomics data sets identified a relatively high number of common metabolites that were decreased after infection with mycobacteria in the human, mouse and zebrafish samples. The comparisons of common metabolites from the humans, mice and zebrafish larvae are shown in the Venn diagram of Fig. 3A. As shown in Fig.3B, the ten common metabolites, methionine, asparagine, cysteine, threonine, serine, tryptophan, leucine, citrulline, ethanolamine and phenylalanine, were significantly decreased in all three sample sets. It should be noted that the overlap between species was limited by the fact that some metabolites that were detected in the human blood samples could not be detected in the mouse or zebrafish samples. This result is true for both the original and normalized values of these ten common metabolites (Fig.5). Therefore these metabolites represent good markers for TB disease activity. Remarkably, five metabolites of this marker set have been previously identified as markers for TB disease in different human populations^3,16^. Three of these markers have also been shown to be predictive for progression of TB. Furthermore, in some cases, such as histidine in the zebrafish samples, a positive correlation was observed, but the threshold of significance was not achieved for inclusion in the overlap set (Fig.3B). In case of zebrafish, this could be the result of the small size of single larva of approximately 0.3 microliter that limits the amount of material analysed^31^. In order to verify the zebrafish data in more detail we therefore undertook an NMR study with larger numbers of pooled zebrafish larvae.

**Fig.5.**
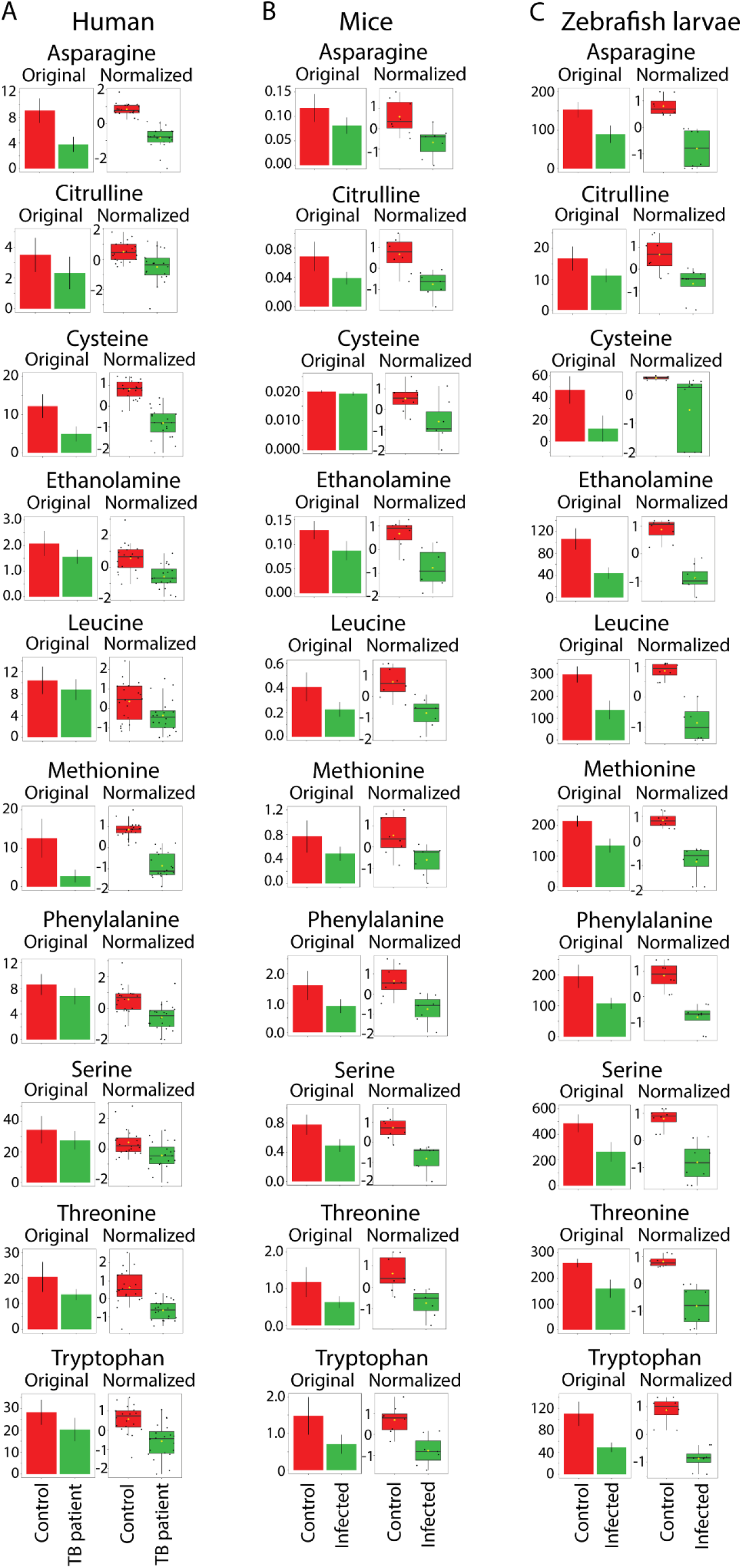
Quantification of the common 10 amino acids that have a decreased concentration in the infected groups of humans, mice and zebrafish. The data for the 10 common metabolites which all display a significantly decreased concentration (p< 0.05) in all the infected groups are shown for **A.** the patients with active TB disease, **B.** mice, and **C.** zebrafish larvae, with the original and normalized values.

### Metabolic profiles in *M.marinum* infected zebrafish larvae measured by nuclear magnetic resonance

In order to confirm the alterations in metabolism in zebrafish after infection with *M.marinum*, ^1^H-nuclear magnetic resonance was used. Metabolic profiles of extracted zebrafish embryos infected for 5 days with the *M.marinum* M strain were obtained by one-dimensional (1-D) and two-dimensional (2-D) ^1^H NMR. Representative 1D ^1^H-NMR spectra from mock-injected control and infected group are shown in figure 6A. 1D ^1^H-NMR of *M.marinum-injected* embryos chemical shift were compared to controls (Fig.6A). The ^1^H-NMR spectra of control and *M.marinum*-injected embryos were investigated by multivariate analysis with PLSDA modeling to probe if control and infected embryos can be discriminated. The PLSDA score plot of the first two principle components explains 38.5% of the total variance, and clustering of the control and infected larvae groups could be observed in the score plot of PLSDA1 vs PLSDA2 (Fig.6B). Comparisons (based on 1-D NMR chemical shifts) to the Human Metabolome Database (HMDB), along with 2-D NMR techniques (i.e., 1H-1H COSY) enabled unambiguous identification and subsequent quantitation of metabolites (Fig.S4).

**Fig.6.**
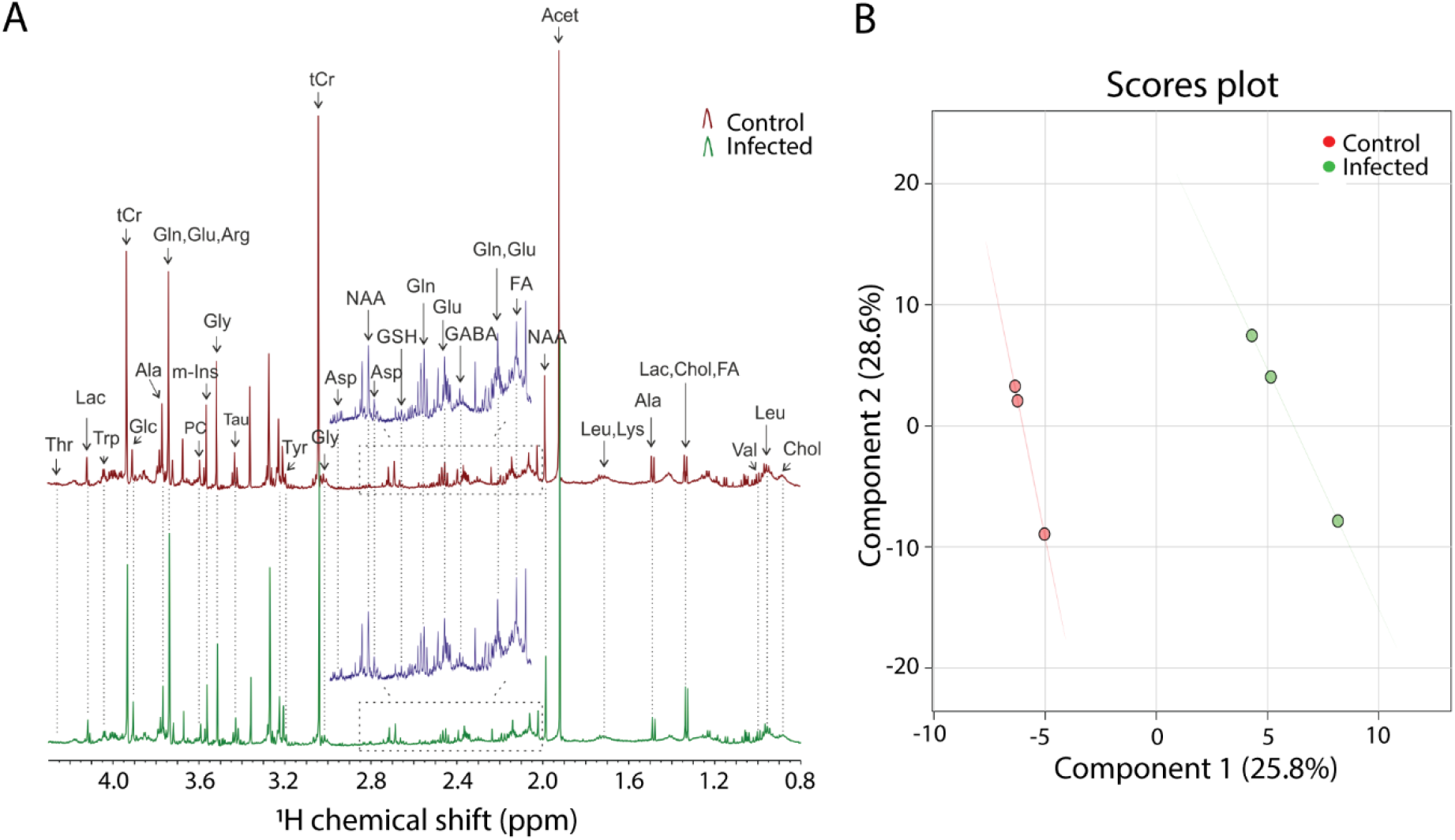
One-dimensional ^1^H NMR spectra of *M.marinum-infected* and uninfected zebrafish larvae. **A**. Zebrafish larvae at 4 hours post fertilization were infected with the *M.marinum* M strain in the yolk and NMR measurements were carried out at 5 days post fertilization. Spectra from chemical shift 0.8 to 4.3 were assigned to specific metabolites. Abbreviations: Thr=threonine, Lac=lactate, Trp=Tryptophan, tCr=total creatine (creatine + phosphocreatine), Glc=Glucose, Ala=alanine, Gln=glutamine, Glu=glutamate, Arg=arginine, PC=phosphocholine, m-Ins=myo-inositol, Gly=glycine, Tau=taurine, Tyr=tyrosine, Asp=aspartate, NAA=N-acetyl aspartate, GSH=glutathione, GABA=gamma-aminobutyric acid, FA=fatty acid, Leu=leucine, Lys=lysine, Val=valine, Chol=cholesterol. **B**. PLSDA shows a clear separation of control and infected groups, n=3, each replicate represents 120 pooled larvae.

### Nuclear magnetic resonance confirms data from mass spectrometry in zebrafish larvae

To compare the consistency of metabolic changes in zebrafish performed by MS and NMR, quantification and comparison of the most abundant metabolites from two approaches between control and *M.marinum*-infected zebrafish embryos are shown in Fig.7. Quantitative analysis of metabolites which were identified in both MS and NMR showed the same trend of changes for all metabolites (Fig.7A, B). The significant changes included 12 decreased metabolites (Fig.7A) that included the entire group of the 10 common biomarker set from the MS analyses. A common biomarker in the human and mouse data set, histidine, which was not significant in the MS analysis in zebrafish, however was significantly decreased in the infected group in the NMR analysis (Fig.7C). Notably, the increased level of putrescine after infection was confirmed by NMR analysis of the zebrafish larvae (Fig.7A, B). NMR could detect additional significantly changed metabolites which were not found in MS, as shown in Fig.7C: lactate, taurine and GABA were reduced in the infected group, while glucose, mannose, and NADH were significantly increased after infection.

**Fig.7.**
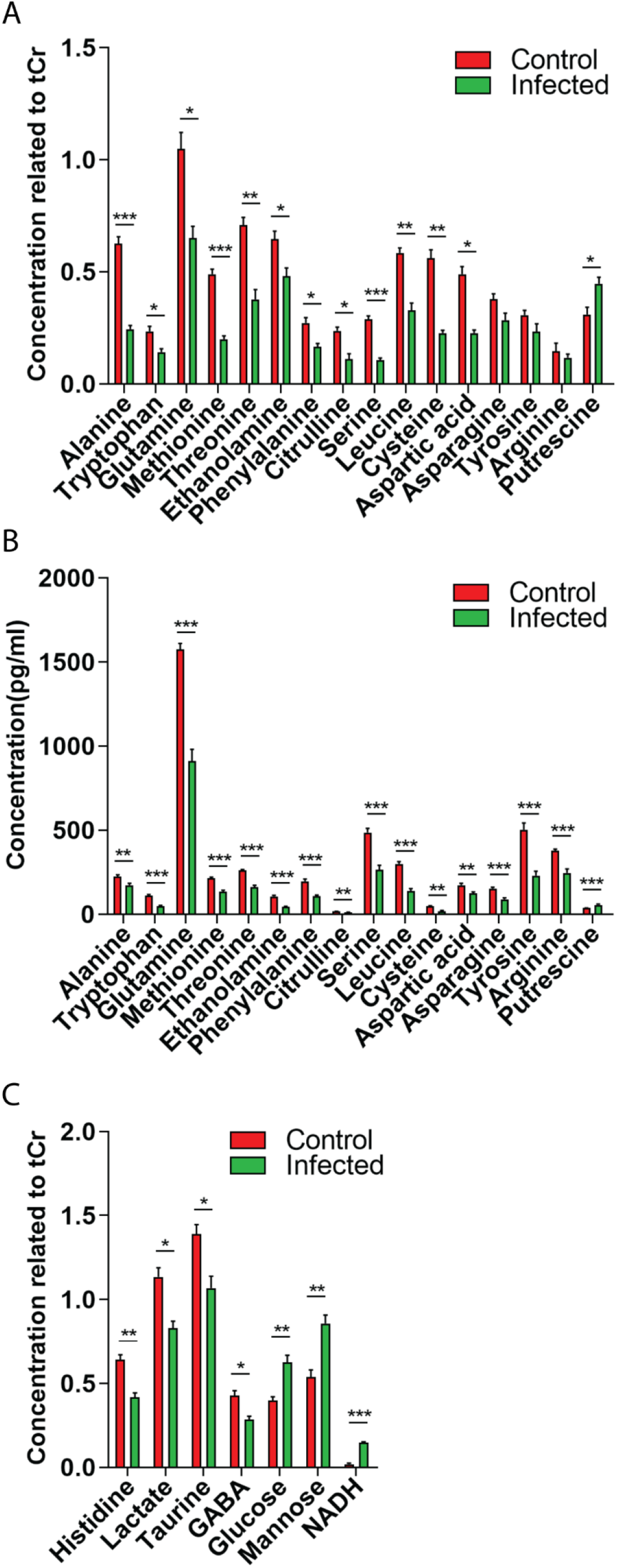
Comparison of the effect of *M.marinum* infection in zebrafish on the metabolic profiles measured by ^1^H NMR measurements and mass spectrometry. **A.** Quantification of metabolite concentrations obtained by NMR spectroscopy. Values are average ± SE of mean (<0.05). tCr represents total creatine. **B.** MS data of the corresponding metabolites shown in Fig.4 and Table S3. **C.** Metabolites which are significantly changed in concentration identified by NMR, but not by MS. Asterisks represent significant p-values obtained by a comparative T-test. \**p*<0.05, \*\**p*<0.01, \*\*\**p*<0.001.

## Discussion

Measurements of metabolites in human TB patients have shown to be a promising method for diagnostic purposes. However, in the absence of knowledge about genetic, physiological and environmental factors that could influence the measurements at different stages of the disease, clinical implications are still uncertain. Our data of metabolic changes in a small contingent of tuberculosis patients in the Netherlands confirm the notion that the observed changes compared to healthy individuals are very common for patients in highly different geographic locations and showed the consistency of our highly-standardized metabolite analysis pipeline^32,33^. In this study we also compared host metabolic changes during human TB disease to infection with *Mtb* in a rodent model and infection with *M.marinum* in a zebrafish using identical pipelines. Our results show highly consistent metabolic changes in all three models. This internal validation of the findings in quite different models enhances the biological broad impact of the findings in the human patients. This is surprising because the analyzed samples of this comparative infection study are in many respects extremely different in that the species represent quite diverse examples from the vertebrate subphylum, that differ for example in metabolic rate, body size, body temperature and examined developmental stages. Since zebrafish larvae are not yet fed during the timeframe of the experiment, but solely use their yolk for nutrition, this shows that difference in food intake are not involved in the observed metabolic changes. Considering these differences, it is striking that in the three studies we found 10 common metabolites that were significantly decreased during infection or disease from a total of 89 metabolites that were detectable with our mass spectrometry platform. These data suggest that this set of metabolites is strongly conservedduring mycobacterial infection and disease in all species.

NMR was used to confirm the mass spectrometry results in the case of zebrafish. These results confirm results from earlier studies that metabolic changes during TB are very significant. Importantly, here we show for the first time that the metabolic changes during TB are highly similar within different vertebrates, at least for the metabolites measured. Apparently, these metabolic changes are so fundamental to mycobacterial infection that they are not affected by the many factors that we mention above. This observation strengthens the biological significance of the finding in human patients, and also supports the potential utility of these metabolites as biomarkers for TB in different trans-species settings, including humans and animal TB, and also diseases caused by non-tuberculosis mycobacteria such as *M.marinum*. In addition, the conservation of these biomarkers in model organisms provides a way to further investigate the mechanism underlying the changes in biomarker levels during TB. Since these biomarkers are common between humans, mice and fish, kinetic analyses that will contribute significantly to pathophysiologic and mechanistic insights can be performed in animal models.

Out of the 10 identified common biomarkers whose levels are decreased in our study, 5 represent amino acids that are essential in human diet^34^. Several of the other decreased biomarkers that are not conserved in all three trans species data sets are also essential amino acids or are derived from these essential amino acids such as tyrosine that is derived from phenylalanine. The decreased levels of these amino acid biomarkers are consistent with earlier reports by Weiner et al, 2012, Weiner et al, 2018, and Vrieling et al, 2019 (Fig.3B). These results indicate that differences in diet or biosynthesis are unlikely to be causing the decreased levels of the identified common group of metabolites. The fact that these biomarkers are also reduced in zebrafish larvae infected with mycobacteria shows an uncoupling of the biomarkers with food intake. One could argue that the decreased levels of these metabolites are the result of consumption of circulating host metabolites by the mycobacteria. This hypothesis seems rather unlikely in the light of the observation that short term treatment with antibiotics did not alter the levels of these metabolites in human patients, despite strongly reducing the number of viable bacilli. It is therefore more likely that the direct cause of the decreased levels of metabolites is the consumption of these metabolites by host cells as a result of inflammation resulting in wasting syndrome. This would be in line with the fact that infection with mycobacteria leads to a metabolic wasting syndrome that can last 6 months or longer after initial antibiotic treatment of the disease^35^. In a previous study, treatment of TB patients with antibiotics indeed lead to recovery of the levels of amino acids after 26 weeks, but this recovery was not yet complete after 8 weeks of treatment^16^.

Currently the mechanisms underlying the strong effects of mycobacterial infection on host metabolism are still poorly understood. Previous studies have suggested that some metabolites might be indicative of changes in particular enzymes involved in host metabolism linked to *Mtb* infection. During TB, an increased level of the enzyme indoleamine 2,3 dioxygenase 1 (IDO1) that converts tryptophan to kynurenine might be responsible for decreased levels of tryptophan and an increased level of kynurenine^3,36^. Our results also indicate a decreased level of tryptophan in the infected state of all three species, but, we have not been able to detect increased levels of kynurenine in the human Groningen contingent and zebrafish samples. Surprisingly, we found a decrease of kynurenine after infection in the mouse experiments (Table S3). Other amino acids that are changed in our data sets such as citrulline, ornithine, arginine and aspartic acid are intermediates in the urea cycle that functions in the conversion of ammonia to urea. The only consistently decreased metabolite of these urea cycle intermediates in all three species is citrulline. Interestingly, previous studies have shown that arginine depletion as a result of arginase activity can be replenished from citrulline in macrophages and T-cells and thereby plays a role in immune defense^37,38^. Although in zebrafish larvae, the T-cells are not yet developed^39^, functional macrophages that can express arginase are present^40^. In addition, zebrafish larvae produce various cytokines that have been found to correlate with decreased amino acid levels in human patients^3^. Therefore, the reduced citrulline levels in zebrafish larvae could indicate a link with innate immune responses. The zebrafish infection model as also used in this study has been recently analyzed using a mathematical model which can predict zebrafish metabolism from gene expression. The results showed that the model predicts reduced histidine metabolism in *M.marinum* infected zebrafish larvae at 4 days post injection^41^. In accordance with the model, a reduction of histidine levels in infected zebrafish larvae was shown to be significant as measured by NMR.

In conclusion, our study provides robust, cross species and validating metabolic marker changes during TB and provides important leads to investigate mechanisms underlying this metabolic reprogramming in animal models. Moreover, further studies should address whether the observed metabolic changes during TB can result in meaningful biomarkers for *Mtb* infection and disease^42^. Animal models can be used to analyse the effect of genetic mutants in key metabolic pathways that could play a role in the observed changes of metabolism. In our future studies we will be interested in genes that play a role at the interface of immune response and metabolism for which knock out mutants are available such as genes in toll-like receptor, leptin^43^ and glucocorticoid receptor^44^ signalling pathways. These studies could not only help elucidating the observed changes in metabolism during infection, but could also lead to the development of new medicines for treatment of TB or wasting syndrome using innovative host-directed therapeutic approaches^45^.

## Methods

### Biological material

Human plasma samples were collected at the specialized UMCG clinical center for TB treatment, Beatrixoord in Haren, Groningen, The Netherlands. All patients were diagnosed with active pulmonary and/or extrapulmonary *M.tuberculosis* infection. Patients were diagnosed with active pulmonary or extrapulmonary TB using mycobacterial culture or PCR of sputum or tissue specimens. For all culture positive specimens, drug sensitivity was determined to guide clinical treatment. All patients improved clinically upon treatment with anti-TB drugs. Patients were admitted to the TB ward for clinical treatment and received a 6 month antibiotic regimen based on Dutch guidelines. Patients originated from various countries, including TB endemic regions. A second blood sample was collected from all individuals 6 weeks after inclusion into the study (referred to as on treatment sample in Fig.S1). In addition, a control group was recruited at the Leiden University Medical Center, comprised of healthy skin test negative individuals. All human sample collection and research was performed in accordance with the guidelines and regulations of the Leiden University Medical School at time of recruitment under protocol P07.48. The age and gender of the patients and healthy persons are given in table S1A and S1B showing that there is a highly unequal distribution of the gender in the patient group. For the mice experiments, two groups of male mice (strain C57BL/6) were obtained from Charles River Laboratories. Male mice were chosen because they have less variation due to the hormonal cycle. Eight male mice nasally-infected with *M.tuberculosis* strain H37Rv at 6 weeks age, and eight mock infected male mice were kept under standard conditions for 8 weeks. The results showed that the mice were systemically infected as shown by plating of bacteria from isolated organ material (Fig.2B and C). One mouse of the group of 8 infected mice had to be killed at an early stage due to unexpected symptoms. There was no significant difference in body weights in the infected groups (Fig.2A). Zebrafish breeding and embryo collection were performed as described previousy^46^. Zebrafish embryos at 2 to 4 hours post fertilization (hpf) were used for *M.marinum* injection. For mass spectrometry analysis we used *M.marinum* strain E11 labelled with a mCherry-expressing pSMT3 vector^26^. For the NMR experiments we used *M.marinum* strain M labelled with mWasabi plasmid pTEC15 vector^22^. *M.marinum* preparation and injection were followed by the protocol of a previous study^26^. Infection levels were checked with fluorescence microscopy (Fig. S3A, B, C and D). One larva per sample (n=8) obtained from different crossings that show batch effects in the clustering was used for LC-MS/MS analysis. Regarding the limitation of NMR sensitivity, 120 larvae per sample (n=3) with *M.marinum* injection or 2% polyvinylpyrrolidone 40 (PVP40) mock injection were used for NMR measurements.

### LC-MS

Human plasma was derived from heparinized tubes. Mice serum was derived of clotted blood tubes, aliquoted, and stored at −80 °C until analysis. A zebrafish embryo has a volume of approximately 0.3 μL^31^. Because of this small volume it is important to dilute the sample as minimal as possible in order to obtain a comprehensive metabolic profile. For this reason the derivatization volumes were downscaled to 4μL:16μL (AccQ:borate) which made the sample 2,5 times less diluted. Although the sample composition didn’t change by this optimization step, the sample was too organic (20% ACN) to give the polar metabolites good chromatography. This problem was solved by diluting the sample four times with water, so the ACN concentration in the sample went to 5%. To compensate for this dilution factor, the sample injection volume was increased with the same factor, so from 1μL injection to 4 μL injection. When the larvae are ready to be analyzed, they are transferred to (1.5mL) centrifuge tubes and are placed in a random order for the washing and quenching by methanol. After homogenization, the proteins were removed from the sample by centrifuging and collecting the supernatant. After collecting the supernatant (75μL), a small portion (5μL) is used for a QC pool which was depending on the batch size aliquoted over 5 centrifuge tubes. The QCs and samples are all placed randomly in the evaporator and derivatized afterwards. After derivatization the samples were diluted with water four times and transferred into vials that were randomly placed in the auto sampler of the LC.

The amine platform covers amino acids and biogenic amines employing an Accq-Tag derivatization strategy adapted from the protocol supplied by Waters^47^. 5.0 μL of each sample was spiked with an internal standard solution. Then proteins were precipitated by the addition of MeOH. The supernatant was taken to dryness in a speedvac. The residue was reconstituted in borate buffer (pH 8.5) with AQC reagent. 1.0 μL of the reaction mixture was injected into the UPLC-MS/MS system. Chromatographic separation was achieved by an Agilent 1290 Infinity II LC System on an Accq-Tag Ultra column (Waters). The UPLC was coupled to electrospray ionization on a triple quadrupole mass spectrometer (AB SCIEX Qtrap 6500). Analytes were detected in the positive ion mode and monitored in Multiple Reaction Monitoring (MRM) using nominal mass resolution. Acquired data were evaluated using MultiQuant Software for Quantitative Analysis (AB SCIEX, Version 3.0.2). The data are expressed as relative response ratios (target area/ISTD area; unit free) using proper internal standards. For analysis of amino acids their ^13^C^15^N-labeled analogs were used. For other metabolites, the closest-eluting internal standard was employed. In-house developed algorithms were applied using the pooled QC samples to compensate for shifts in the sensitivity of the mass spectrometer over the batches. After quality control correction, metabolite targets complied with the acceptance criteria of RSDqc <15%. Using this platform we were able to identify maximally 89 metabolites in blood samples from humans, mice and total zebrafish larvae. Not all these metabolites could be detected in each of the species.

### MS data analysis

Data was analysed using the software package MetaboAnalyst 4.0^48^. MetaboAnalyst offers the possibility to provide automated data reports which we used for archiving data sets. In brief, default settings were used with log transformation and auto scaling of the data for normalisation (Fig. 5). PLSDA was performed with PLS regression using the plsr function provided by R pls package. The classification and cross-validation are performed using the corresponding wrapper function offered by the caret package^49^. Naming of the metabolites is based on reference compounds using standard nomenclature of the human metabolome database (http://www.hmdb.ca/). We have categorized cystine and cysteine as one metabolite.

### NMR spectroscopy

For NMR spectroscopy, metabolites from zebrafish larvae were extracted based on an adapted protocol from a previous study^50^. In brief, 120 zebrafish larvae per sample were crashed in the mixture of methanol: water (1:1, v/v). Subsequently, 1ml chloroform was added. The mixture was sonicated for 15 minutes and then centrifuged at 5000rpm for 5 minutes. After centrifugation, two layers were formed: the upper layer is methanol and water. The dried methanol:water layer with metabolites was dissolved in 1ml of 100mM deuterated phosphate buffer (KD2PO4, PH=7.0) containing 0.02% trimethyl-silylpropanoic acid (TSP) as internal standard and subsequently filtered with Millipore filter (Millex-HV 0.45-lmFilterUnit). Metabolites in zebrafish larvae were measured with a Bruker DMX 600MHz NMR spectrometer at 4°C equipped with a 5mm inverse triple high-resolution probe with an actively shielded gradient coil. The ^1^H NMR spectra were accumulated with 65,000 data points, a 2-s relaxation delay, a sweep width of 12.4 kHz, and 256 scans which were required to obtain a satisfactory signal-to-noise ratio. Two-dimensional homonuclear ^1^H-^1^H experiments were carried out using the chemical shift correlated spectroscopic (COSY) sequence. The parameters used for COSY were 2048 data points collected in the t2 domain over the spectral width of 4k, 512 t1 increments were collected with 16 transients, relaxation delay 2 sec, acquisition time 116 msec, and pre-saturated water resonance during relaxation delay. The resulting data were zero filled with 2048 data points, and were weighted with sine bell window functions in both dimensions prior to Fourier Transformation. The chemical shifts are relative to TSP.

### NMR analysis

NMR analysis was performed based on an adjusted version from previous studies^33,51^. In brief, the one-dimensional ^1^H NMR spectra obtained from both in control and infection group were corrected for baseline and phase shifts using the MestReNova software version 11.0 (Mestrelab Research S.L., Santiago de Compostela, Spain). The spectra were then subdivided in the range between 0 and 10 ppm into buckets of 0.04 ppm. The region of 4.20-6.00 ppm was excluded from the analysis to remove the water peak. The resulting data matrix was exported into Microsoft office excel (Microsoft Corporation). This was then imported into MetaboAnalyst 4.0 for PLSDA. Correlation coefficients with p < 0.05 were considered statistically significant. Quantification of metabolites was performed using Chenomx NMR Suite 8.3, which allowed for qualitative and quantitative analysis of an NMR spectrum by fitting spectral signatures from an HMDB database to the spectrum. Metabolite concentrations were subsequently calculated as ratio to tCr. Statistical analysis (t-tests) of the NMR quantification results was performed with GraphPad Prism 8 and p-values smaller than 0.05 were considered significant.

## Data availability

All data underlying this study are included within the manuscript, supplementary material and supporting information file.

## Acknowledgements

We thank Dr. Slavik Koval for assistance with the MetaboAnalyst program.

## Ethical licenses and consent of patients for publication

Zebrafish lines were handled in accordance with the local animal welfare regulations and maintained according to standard protocols (https://zfin.org). This local regulation serves as the implementation of Guidelines on the protection of experimental animals by the Council of Europe, Directive 86/609/EEC, which allows zebrafish embryos to be used up to the moment of free-living (5 days after fertilization). Since embryos used in this study were no more than 5 days old, no license is required by Council of Europe (1986), Directive 86/609/EEC or the Leiden University ethics committee. The mice experiments were performed under ethical license number DEC 14080 (10-07-2014) of Leiden University. The study with human patients material was approved by the Ethical Committee of the Faculty of Medicine, Leiden University Medical Center, and written informed consent was voluntarily signed by all patients and control subjects registered under protocol P07.48 (March 17, 2008).

## Author contributions

R.J.R. and K.E. performed mass spectrometry experiments, Y.D., performed NMR experiments, R.J.R., Y.D., A.A. and H.P.S. bioinformatic analysis, Y.D., W.J.V., M.C.H., S.v.E and R.M.J. performed animal experiments, S.A.J., M.C.H. and W.J.V. obtained samples and ethical permission, A.C.H., T.H., A.A., T.H.M.O. and H.P.S. supervised technical facilities, A.A., S.A.J., M.C.H.,T.H.M.O. and H.P.S. supervised research, Y.D. and H.P.S. wrote first draft of manuscript and made figures and tables, R.M.J., H.P.S., S.A.J., R.J.R., T.H., M.C.H., T.H.M.O. and A.A. conceived the study, H.P.S., R.J.R., Y.D., R.M.J., W.J.V., S.A.J., M.C.H., and A.A. planned experiments and interpreted results, all authors reviewed the manuscript.

