## Supplemental files for "Tuberculosis causes highly conserved metabolic changes in human patients, mycobacteria-infected mice and zebrafish larvae"

### Supplementary Figure S1

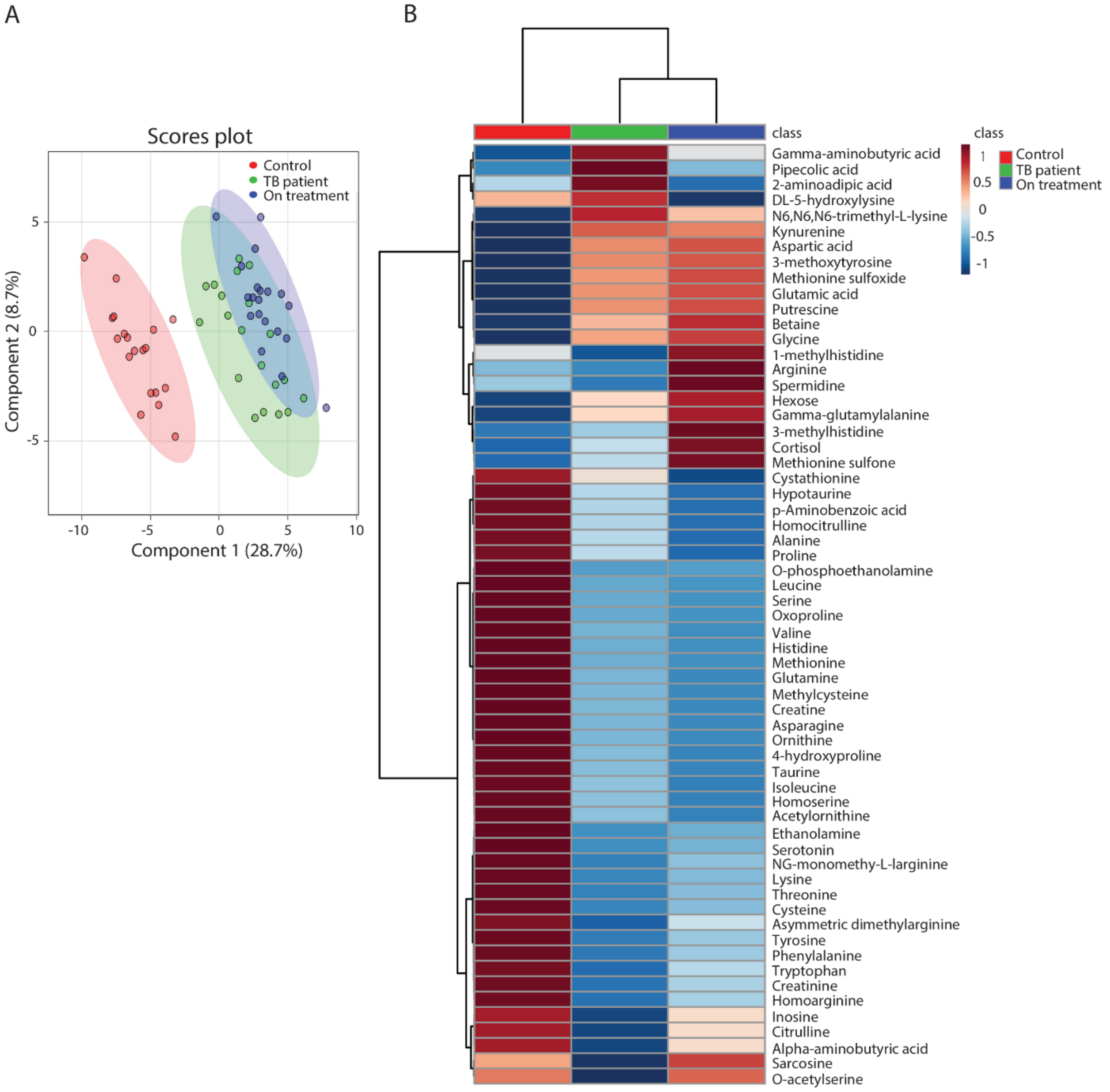


**Supplementary Figure S1. Partial least squares discriminant analysis and heat map of metabolomic profiles from healthy control, TB patient, and on treatment groups. A.** Analysis of blood of the healthy control group (Control), patients with active TB disease (TB patients), and the same patients treated for 6 weeks with antibiotics (On treatment), n=20. **B**. Heat map of all metabolites of blood of the healthy control group (Control), TB infected patients (TB patient) and the same patients treated for 6 weeks with antibiotics (On treatment)

### Supplementary Figure S2

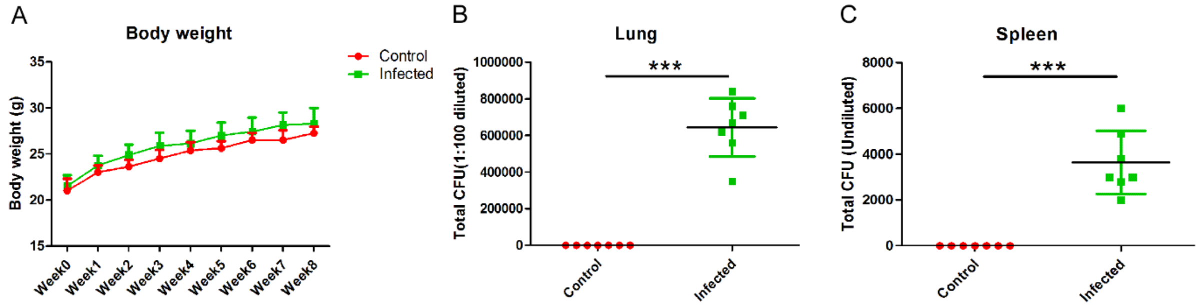


**Supplementary Figure S2. Body weight and total cfu of isolated organs in mice. A.** Body weight of control and infected mice from week 0 to week 8. **B.** Total cfu (1:100 diluted) of isolated lung from control and infected mice after 8 weeks of systemically infection with *Mtb*. **C.** Total cfu (Undiluted) of isolated spleen from control and infected mice after 8 weeks of systemically infection with *M.tb*. ***$p$<0.001. Abbreviation: cfu, colony forming unit.

### Supplementary Figure S3

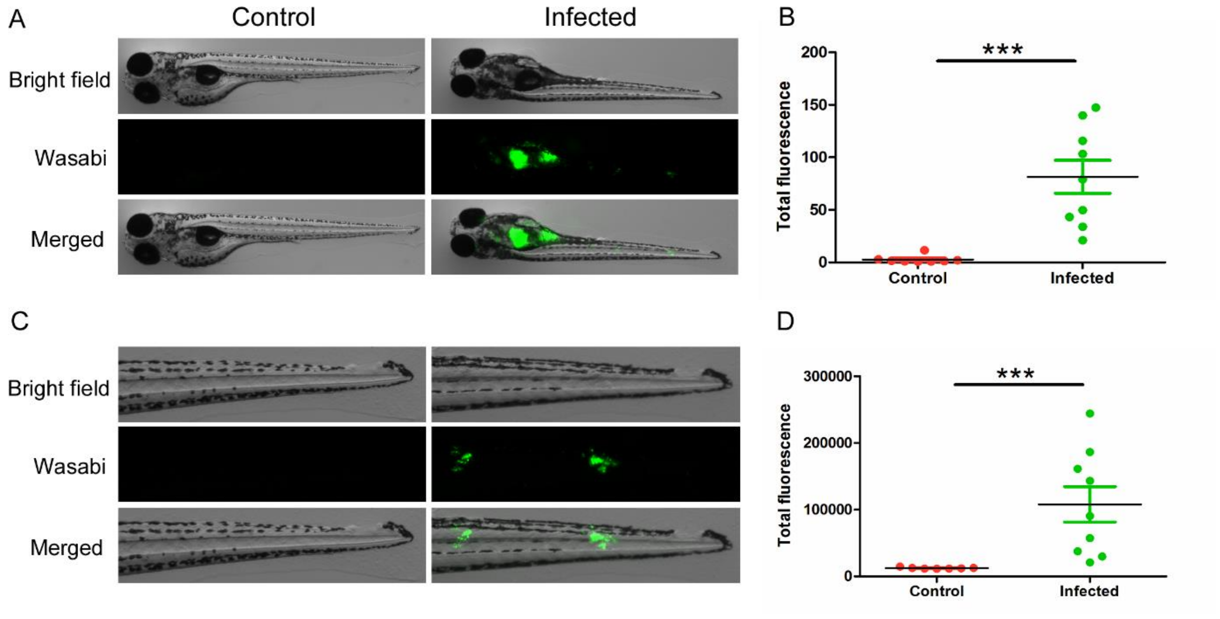


**Supplementary Figure S3. Representative images and quantification of *M.m* infection in zebrafish larvae. A.** The representative images of whole larva from control and infected group. **B.** Quantification of total fluorescence of wasabi from whole larva in two groups. **C.** The representative images of tail part from control and infected group. **D.** Quantification of total fluorescence of wasabi from tail part in two groups.

### Supplementary Figure S4

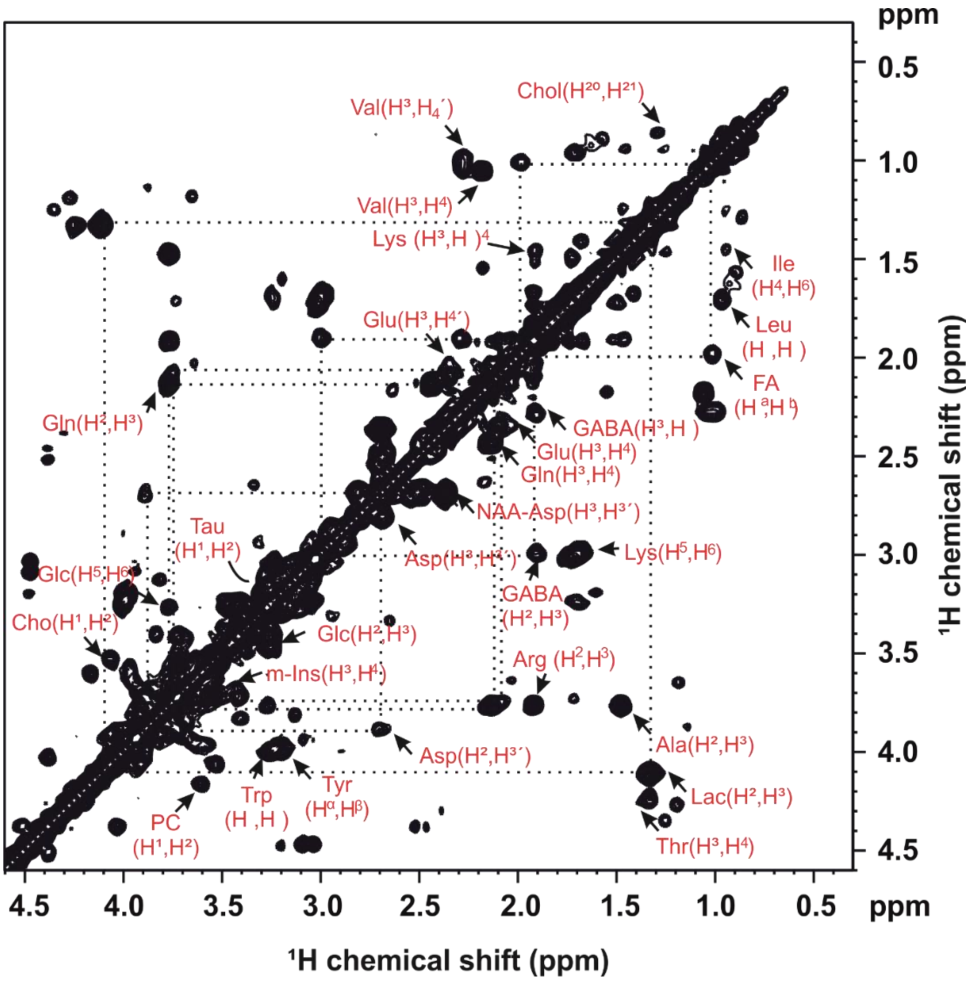


**Supplementary Figure S4. Representative high resolution 2D $^1$H-$^1$H homonuclear correlation spectrum in control zebrafish larvae.** The extracted metabolites of zebrafish larvae obtained using the chemical shift correlated spectroscopic (COSY) sequence.

### Supplementary Table S1.A

| Gender and Age | Control<br>n=20 | TB patient<br>n=20 |
|---|---|---|
| Gender(male/female) | 7/13 | 18/2 |
| Age (years) | 34.5+12.4 | 32.9+14.8 |

### Supplementary Table S1.B

| Sample | Label | Gender | Age |
|---|---|---|---|
| C5_1 | Control | female | 33 |
| C5_3 | Control | female | 33 |
| C5_7 | Control | male | 25 |
| C5_10 | Control | female | 41 |

| | | | |
|---|---|---|---|
| C5_13 | Control | female | 42 |
| C5_14 | Control | female | 26 |
| C5_15 | Control | female | 31 |
| C5_18 | Control | female | 44 |
| C5_20 | Control | male | 18 |
| C5_21 | Control | female | 54 |
| C5_24 | Control | female | 22 |
| C5_25 | Control | female | 22 |
| C5_27 | Control | male | 69 |
| C5_28 | Control | female | 23 |
| C5_29 | Control | male | 36 |
| C5_30 | Control | male | 32 |
| C5_32 | Control | female | 43 |
| C5_33 | Control | male | 33 |
| C5_34 | Control | male | 41 |
| C5_38 | Control | female | 22 |
| TR001 | TB patient | male | 22 |
| TR002 | TB patient | male | 31 |
| TR003 | TB patient | male | 27 |
| TR004 | TB patient | male | 24 |
| TR005 | TB patient | male | 44 |
| TR006 | TB patient | male | 31 |
| TR007 | TB patient | male | 34 |
| TR008 | TB patient | male | 48 |
| TR009 | TB patient | male | 41 |
| TR010 | TB patient | male | 26 |
| TR011 | TB patient | male | 29 |
| TR012 | TB patient | male | 23 |
| TR013 | TB patient | male | 21 |
| TR014 | TB patient | male | 29 |
| TR016 | TB patient | female | 85 |
| TR017 | TB patient | male | 43 |
| TR018 | TB patient | female | 32 |
| TR019 | TB patient | male | 24 |
| TR020 | TB patient | male | 17 |
| TR021 | TB patient | male | 26 |

**Supplementary Table S1. Gender and age information in healthy people and TB patients.** **A.** Summarized gender and age information from two groups. **B.** Individual gender and age information from two groups.

**Supplementary Table S2**

| Metabolites | HMDB identifier | Human FDR | Human Ratio |
|---|---|---|---|
| Methionine | HMDB00696 | 1,01E-11 | 0.22 |
| Methionine sulfoxide | HMDB02005 | 1,47E-11 | 5.78 |
| Serotonin | HMDB00259 | 5,35E-11 | 0.003 |
| Asparagine | HMDB00168 | 5,35E-11 | 0.42 |
| Cysteine | HMDB00574 | 2,47E-09 | 0.40 |
| Methylcysteine | HMDB02108 | 4,05E-08 | 0.48 |

| | | | |
|---|---|---|---|
| Hypotaurine | HMDB00965 | 9,09E-08 | 0.23 |
| Aspartic acid | HMDB00191 | 4,04E-07 | 1.93 |
| Glutamic acid | HMDB00148 | 1,04E-06 | 1.97 |
| Glutamine | HMDB00641 | 1,21E-06 | 0.33 |
| Lysine | HMDB00182 | 1,90E-06 | 0.73 |
| O-phosphoethanolamine | HMDB00224 | 1,59E-05 | 0.0003 |
| Threonine | HMDB00167 | 2,94E-05 | 0.66 |
| Taurine | HMDB00251 | 5,84E-05 | 0.60 |
| NG-Monomethy-L-arginine | HMDB29416 | 1,72E-04 | 0.57 |
| Ethanolamine | HMDB00149 | 2,50E-04 | 0.75 |
| Tryptophan | HMDB00929 | 2,55E-04 | 0.72 |
| Histidine | HMDB00177 | 3,72E-04 | 0.46 |
| Phenylalanine | HMDB00159 | 4,16E-04 | 0.79 |
| Homoarginine | HMDB00670 | 7,77E-04 | 0.62 |
| Alpha-aminobutyric acid | HMDB00452 | 9,00E-04 | 0.75 |
| Putrescine | HMDB01414 | 1,51E-03 | 3.18 |
| Citrulline | HMDB00904 | 2,00E-03 | 0.67 |
| Homoserine | HMDB00719 | 2,46E-03 | 0.75 |
| Serine | HMDB00187 | 8,84E-03 | 0.80 |
| Asymmetric dimethylarginine | HMDB01539 | 1,16E-02 | 0.79 |
| Inosine | HMDB00195 | 1,46E-02 | 0.16 |
| Gamma-aminobutyric acid | HMDB00112 | 1,85E-02 | 1.47 |
| p-Aminobenzoic acid | HMDB01392 | 2,30E-02 | 0.79 |
| Creatine | HMDB00064 | 3,35E-02 | 0.67 |
| Leucine | HMDB00687 | 3,50E-02 | 0.84 |

**Supplementary Table S2. Ratio of metabolite quantities in blood of TB patients compared to the control group.** Ratios of metabolite quantities in human blood samples. The levels of 31 metabolites are significantly altered in TB disease compared to healthy people.

**Supplementary Table S3**

| Metabolites | HMDB identifier | Mice FDR | Mice ratio |
|---|---|---|---|
| Sarcosine | HMDB00271 | 4,63E-05 | 0.36 |
| Ornithine | HMDB00214 | 1,38E-04 | 0.52 |
| Proline | HMDB00162 | 1,92E-04 | 0.53 |
| Serine | HMDB00187 | 3,07E-04 | 0.63 |
| 2-aminoadipic acid | HMDB00510 | 5,80E-04 | 0.40 |
| Glycine | HMDB00123 | 5,90E-04 | 0.63 |
| Gamma-glutamylalanine | HMDB06248 | 8,43E-04 | 0.47 |
| Histidine | HMDB00177 | 9,87E-04 | 0.66 |
| Homoserine | HMDB00719 | 1,21E-03 | 0.59 |
| Tryptophan | HMDB00929 | 1,23E-03 | 0.48 |
| Methionine sulfoxide | HMDB02005 | 1,28E-03 | 0.51 |
| Ethanolamine | HMDB00149 | 1,31E-03 | 0.67 |
| Leucine | HMDB00687 | 1,58E-03 | 0.55 |
| Alanine | HMDB00161 | 1,69E-03 | 0.61 |
| Citrulline | HMDB00904 | 2,11E-03 | 0.57 |
| Phenylalanine | HMDB00159 | 2,62E-03 | 0.56 |
| Threonine | HMDB00167 | 2,78E-03 | 0.54 |
| Isoleucine | HMDB00172 | 2,94E-03 | 0.57 |

| | | | |
|---|---|---|---|
| Arginine | HMDB00517 | 3,15E-03 | 0.61 |
| Kynurenine | HMDB00183 | 3,47E-03 | 0.55 |
| Tyrosine | HMDB00158 | 3,89E-03 | 0.52 |
| Alpha-aminobutyric acid | HMDB00452 | 4,11E-03 | 0.65 |
| Methyldopa | HMDB11754 | 4,30E-03 | 0.52 |
| Gamma-aminobutyric acid | HMDB00112 | 4,38E-03 | 0.57 |
| Valine | HMDB00883 | 5,51E-03 | 0.54 |
| Lysine | HMDB00182 | 5,77E-03 | 0.59 |
| Asparagine | HMDB00168 | 9,04E-03 | 0.69 |
| Glutamic acid | HMDB00148 | 9,68E-03 | 0.64 |
| Methionine | HMDB00696 | 1,79E-02 | 0.63 |
| Cysteine | HMDB00192 | 2,36E-02 | 0.97 |
| Spermidine | HMDB01257 | 2,63E-02 | 0.49 |

**Supplementary Table S3. Ratio of metabolite quantities in blood of *Mtb*-infected mice compared to the control group.** Ratios of metabolite quantities in mice blood samples. The levels of 31 metabolites are significantly altered in the infected compared to the control group.

**Supplementary Table S4**

| Metabolites | HMDB identifier | ZF FDR | ZF ratio |
|---|---|---|---|
| Ethanolamine | HMDB00149 | 9,70E-07 | 0.41 |
| Valine | HMDB00883 | 1,16E-06 | 0.48 |
| Tryptophan | HMDB00929 | 1,38E-06 | 0.45 |
| Isoleucine | HMDB00172 | 1,79E-06 | 0.42 |
| Ornithine | HMDB00214 | 2,01E-06 | 0.58 |
| Leucine | HMDB00687 | 4,97E-06 | 0.46 |
| Glutamine | HMDB00641 | 1,10E-05 | 0.58 |
| Methionine | HMDB00696 | 1,15E-05 | 0.63 |
| Hydroxyproline | HMDB06055 | 2,05E-05 | 0.46 |
| Phenylalanine | HMDB00159 | 2,16E-05 | 0.55 |
| Threonine | HMDB00167 | 2,71E-05 | 0.62 |
| Proline | HMDB00162 | 6,02E-05 | 0.67 |
| Serine | HMDB00187 | 7,26E-05 | 0.54 |
| Tyrosine | HMDB00158 | 8,11E-05 | 0.45 |
| Asparagine | HMDB00168 | 8,31E-05 | 0.58 |
| Putrescine | HMDB01414 | 1,08E-04 | 1.52 |
| Gamma-glutamylalanine | HMDB06248 | 2,60E-04 | 0.41 |
| Glycine | HMDB00123 | 3,43E-04 | 0.63 |
| Arginine | HMDB00517 | 5,34E-04 | 0.65 |
| Asymmetric dimethylarginine | HMDB01539 | 7,94E-04 | 0.54 |
| Aspartic acid | HMDB00191 | 3,01E-03 | 0.72 |
| Citrulline | HMDB00904 | 3,68E-03 | 0.67 |
| Methylcysteine | HMDB02108 | 4,04E-03 | 0.48 |
| Alanine | HMDB00161 | 4,35E-03 | 0.77 |
| Symmetric dimethylarginine | HMDB03334 | 6,04E-03 | 0.46 |
| Glutathione | HMDB00125 | 6,18E-03 | 1.46 |
| 3-methoxytyrosine | HMDB01434 | 7,16E-03 | 0.71 |
| 5-hydroxytryptophan | HMDB00472 | 8,78E-03 | 0.28 |
| Beta-alanine | HMDB00056 | 9,06E-03 | 1.23 |
| Glutathione disulfide | HMDB03337 | 1,22E-02 | 0.78 |

| | | | |
|---|---|---|---|
| Aminoadipic acid | HMDB00510 | 1,37E-02 | 0.78 |
| Cysteine | HMDB00192 | 2,25E-02 | 0.25 |
| O-phosphoethanolamine | HMDB00224 | 2,32E-02 | 1.17 |
| 5-hydroxylysine | HMDB00450 | 3,52E-02 | 1.49 |
| N6,N6,N6-trimethyl-L-lysine | HMDB01325 | 3,59E-02 | 0.62 |

**Supplementary Table S4. Ratio of metabolite quantities in *M.marinum*-infected zebrafish larvae versus control group obtained by MS.** The concentration of 35 metabolites that are significantly changed in the mycobacterial infected group compared to the control group. ZF ratio: zebrafish larvae with *M.marinum* strain E11 infection compared with control.
